# Molecular Diversity and Recombination Patterns of the ORF7 (Nucleocapsid) Gene in *Betaarterivirus americense* Variants Circulating from Lima, Peru

**DOI:** 10.64898/2026.02.24.707833

**Authors:** Rony Cotaquispe Nalvarte

## Abstract

Porcine reproductive and respiratory syndrome (PRRS) is a globally distributed disease caused by *Betaarterivirus europensis* (PRRSV-1) and *Betaarterivirus americense* (PRRSV-2). Its clinical presentation ranges from subclinical infection to severe disease, depending on viral evolution and the emergence of novel variants. The aim of this study was to characterize the genetic diversity and identify recombination events in the ORF7 (nucleocapsid, N) gene of ten PRRSV-2 strains circulating in pig farms in Lima, Peru. Bioinformatic analyses were performed using DNAMAN v10.0, MEGA 6, BepiPred-2.0, DnaSP v6, and RDP v4.101. Phylogenetic analysis revealed two well-defined lineages: eight strains clustered within lineage 1A (NADC34-like), and two within lineage 5A (VR2332-like), demonstrating the co-circulation of genetically distinct variants in the region. Comparative sequence analysis identified significant amino acid substitutions in eight strains (15, 16, 17, 20, 21, 22, 23, and 24), with strain 24 being the most divergent, accumulating multiple substitutions, including T81I, R109S, I115F, R116S, and A119K within the C-terminal region encompassing antigenic domains I–V. B-cell epitope prediction using BepiPred-2.0 identified six epitope patterns (A–F) comprising nine potential B-cell epitope regions (positions 5–19, 33–72, 33–73, 84–85, 87–98, 84–98, 87–97, 84, and 119). Patterns B, E, and F exhibited four to five predicted epitope sites and corresponded to strains 21, 22, 23, and 24. Recombination analysis using RDP v4.101 detected a statistically robust recombination event in strain 18_montana2020-R (lineage 5A), with strain 24_montana2020-WT (lineage 1A) identified as the putative major parent (100% similarity) and the vaccine-like VR2332 strain (lineage 5A) as the minor parent (99.3% similarity). Secondary evidence of the same recombination event was observed in strain 19_montana2020-R. Genetic diversity analysis of the ORF7 gene identified 50 polymorphic nucleotide sites and 52 mutations. Overall, these findings demonstrate substantial genetic variability in the ORF7 gene of PRRSV-2 circulating in Lima, Peru, characterized by lineage co-circulation and inter-lineage recombination. Continuous molecular surveillance is warranted to monitor viral evolution, assess potential antigenic implications, and support effective PRRS control strategies in the Peruvian swine industry.

## INTRODUCTION

Porcine reproductive and respiratory syndrome virus (PRRSV) remains a persistent threat to global swine production, causing substantial economic losses estimated at over USD 1.2 billion annually in the United States and Europe alone [10]. PRRSV is endemic in most pig-producing countries and continues to pose significant challenges to animal health and the swine industry worldwide. It is an enveloped, positive-sense, single-stranded RNA virus approximately 15 kb in length, containing at least 11 open reading frames (ORFs). ORF1a and ORF1b encode the nonstructural polyproteins pp1a and pp1ab, respectively; structural ORFs 2–6 and ORF5a encode envelope-associated proteins, whereas ORF7 encodes the nucleocapsid (N) protein, the most abundant viral protein in infected cells [1,25].

The ORF7 gene is approximately 372 bp in length and encodes a ∼15 kDa protein lacking N-glycosylation sites. It contains at least five immunologically relevant epitopes located within regions 30–52, 37–52, and 69–123, as well as two classical nuclear localization signals (NLS) and a conserved region known as the superimposed signal for nuclear localization (SSLNu) [33,34,37], underscoring its structural and immunological relevance. Due to its relative conservation combined with sufficient variability to discriminate viral variants, ORF7 is widely used as a molecular target for conventional PCR detection and evolutionary analyses [3,37].

Taxonomically, PRRSV is currently classified into two distinct species according to the International Committee on Taxonomy of Viruses (ICTV): Betaarterivirus europensis (PRRSV-1) and Betaarterivirus americense (PRRSV-2), which share approximately 60% genomic similarity. Phylogenetic classification, primarily based on ORF5 sequences, identifies multiple lineages and sublineages, reflecting the extensive genetic diversity of PRRSV [4,11,35].

In addition to the high mutation rate characteristic of RNA viruses, genetic recombination plays a critical role in PRRSV evolution. Recombination occurs when the RNA-dependent RNA polymerase switches templates during replication, generating mosaic genomes composed of segments derived from different parental strains. This process has been extensively documented and is recognized as a major evolutionary driver of PRRSV-2, often resulting in phylogenetic incongruence between genomic regions such as ORF5 and whole-genome sequences [23].

In the present study, multiple bioinformatic tools were employed to detect and validate recombination events within the ORF7 gene. Recombination Detection Program (RDP) v4.101 [20,23] was used to identify recombination signals through integrated algorithms (RDP, GENECONV, BootScan, MaxChi, Chimaera, SiScan, and 3Seq), increasing robustness by considering only events supported by multiple methods. Genetic diversity and polymorphic site analyses were conducted using DnaSP v6 [26], and phylogenetic analyses were performed using MEGA [20], as previously described by Guo et al. [7]. Our findings support that, beyond the accumulation of point mutations, recombination contributes to genetic diversification even within relatively conserved genomic regions such as ORF7. These processes have important implications for molecular surveillance, phylogenetic characterization, and the development of effective PRRSV control strategies.

## MATERIALS AND METHODS

### Biological Material

A total of ten nucleotide sequences obtained from PRRSV-2 strains previously isolated and characterized by Cotaquispe [3] were included in this study.

### RNA Extraction and cDNA Synthesis

Viral RNA extraction and complementary DNA (cDNA) synthesis of *Betaarterivirus americense* were performed as previously described by Cotaquispe [3]. Reverse transcription was conducted using a Biometra TOne thermocycler (Analytik Jena, Germany) following the manufacturer’s instructions for the OneScript® Hot cDNA Synthesis Kit (ABM®, Canada). The first reaction mixture (14.5 µL total volume) contained 5 µL of viral RNA (40 ng/µL), 1 µL of random primers or gene-specific reverse primers, 1 µL of 10 mM dNTP mix, and 7.5 µL of RNase-free water. The mixture was preheated at 65°C for 5 min, chilled on ice for 1 min, and briefly centrifuged. A second mixture (5.5 µL) containing 4 µL of 5× RT buffer, 0.5 µL RNaseOFF Ribonuclease Inhibitor (40 U/µL), and 1 µL OneScript RTase (200 U/µL) was added to the first reaction and gently mixed. Reverse transcription was performed under the following conditions: primer annealing at 25°C for 10 min (for random primers only), cDNA synthesis at 42°C for 50 min, and enzyme inactivation at 85°C for 5 min. The synthesized cDNA was stored at −80°C until further analysis.

### Conventional PCR Amplification of the ORF7 (N) Gene

PCR amplification of the ORF7 gene encoding the nucleocapsid (N) protein was performed using the Biometra TOne thermocycler. Each reaction (20 µL total volume) contained 3 µL of cDNA (40 ng/µL), 10 µL of 2× PCR HotStart Master Mix (ABM®, Canada), 5 µL nuclease-free water, and 1 µL (10 pmol/µL) of each primer: SKJ-F: 5^′^-GGACCGGGGAAGAAGAACAA-3^′^; SKJ-R: 5^′^-AGTGCCGTTCAC CACTCATT-3^′^. These primers were designed in-house based on the ORF7 sequence and validated for sensitivity, specificity, limit of detection, repeatability, and robustness. Thermal cycling conditions consisted of an initial denaturation at 94°C for 5 min; 45 cycles of 94°C for 45 s, 52°C for 60 s, and 72°C for 45 s; followed by a final extension at 72°C for 10 min. The expected amplicon size was 327 bp, corresponding to positions 14,473 to 14,842 of the reference ORF7 genomic region.

### Gel Electrophoresis, Purification, and Sequencing

PCR products were resolved by horizontal gel electrophoresis using a multiSUB™ Choice system (Cleaver Scientific, UK) with 1% molecular-grade agarose (SCL AG500, Cleaver Scientific). Gels were stained with Safe-Green™ DNA dye (ABM®, Canada) and visualized using a UVP UVsolo Touch imaging system (Analytik Jena, Germany). A 100 bp DNA ladder (50 bp–1.5 kb, ABM®) was used as a molecular size marker. Bands of the expected size were excised and purified using the innuPREP Nucleic Acid Purification Kit (Analytik Jena, Germany). Purified amplicons were sequenced bidirectionally by Macrogen Inc. (Seoul, South Korea) using the BigDye® Terminator v3.1 Cycle Sequencing Kit (Applied Biosystems, USA) on an ABI 3130XL Genetic Analyzer (Applied Biosystems).

### Phylogenetic Analysis of the ORF7-Encoded N Protein

The ten chromatograms were edited and assembled using Chromas Lite v2.6.6 (Technelysium Pty Ltd, Australia) (ChromasPro | Technelysium Pty Ltd). Sequences were aligned with 42 reference sequences retrieved from GenBank, representing globally distributed pathogenic strains [12,28]. Phylogenetic relationships were inferred using the maximum likelihood (ML) method based on the Jones–Taylor–Thornton (JTT) amino acid substitution model. Branch lengths were measured as the number of substitutions per site. The final dataset included 131 amino acid positions. Bootstrap support values were calculated using 1,000 replicates. Evolutionary analyses were performed using DNAMAN v10.0 (Lynnon Biosoft, Canada) (https://www.lynnon.com/download/) [31] and MEGA version 6 (https://www.megasoftware.net/) [29].

### Analysis of Antigenic Domains and Linear B-Cell Epitopes

Antigenic domain mapping of the N protein was conducted using MEGA 6.0 [7, 29]. The nucleocapsid protein of *Betaarterivirus americense* consists of 123 amino acids, whereas B. europensis contains 128 amino acids. Structural features analyzed included: N-terminal and C-terminal regions, Nuclear localization signal 1 (NLS-1; aa 10–13), Nuclear localization signal 2 (NLS-2; aa 41–47), Nucleolar localization signal (NoLS; aa 41–72), Conserved cysteine residue at position 23 (Cys23), implicated in intermolecular disulfide bond formation and N protein homodimerization, Serine residue at position 120 (Ser120), a putative phosphorylation site [21, 34]. Five antigenic domains were evaluated: Domain I (aa 30–52), Domain II (aa 37–52), Domain III (aa 52–69), Domain IV (aa 69–112), and Domain V (aa 112–123) [5,8,15,21,28,36]. Amino acid substitutions within immunologically relevant domains were identified, and putative linear B-cell epitopes were predicted using the BepiPred-2.0 server (IEDB Analysis Resource) (http://tools.iedb.org/main/bcell/), based on a 123-amino acid ORF7 protein sequence. Predictions were performed using a Random Forest algorithm trained on experimentally validated epitope datasets [2,13,28].

### Genetic Diversity and Recombination Analysis

Genetic diversity indices were calculated using DnaSP v6 [26]. The ORF7 sequence dataset was initially screened for recombination signals. Recombination detection was performed using seven algorithms implemented in RDP v4.101 [20]: RDP, GENECONV, BootScan, MaxChi, Chimaera, SiScan, and 3Seq [6,16–18,22,24,27]. Default parameters were applied (http://web.cbio.uct.ac.za/~darren/rdp.html). Recombination events were considered reliable only when supported by at least four independent methods with p-values < 0.05. Given the relatively short length of ORF7 (∼372 bp), a more stringent threshold (p < 0.025) with Bonferroni correction for multiple comparisons was applied to reduce false-positive signals. A sliding window size of 20–30 nucleotides with a step size of 5 nucleotides was used to improve detection resolution in this compact genomic region [12,32].

## RESULTS

### Amplification and Sequencing of the ORF7 Gene

Samples were analyzed using an in-house conventional RT-PCR assay targeting the ORF7 gene encoding the nucleocapsid (N) protein. Primers were validated and standardized using type 2 (North American) vaccine strain Nebraska Prime Pac® PRRS and type 1 (European) VP-046 SUIPRABAC® PRRS as positive controls. Successful amplification of a 327 bp fragment corresponding to the ORF7 gene was obtained (Figure 1). Bidirectional sequencing of all amplicons confirmed partial ORF7 gene sequences comprising 369 nucleotides, encoding the complete 123-amino-acid nucleocapsid (N) protein (Figure 3).

**Figure 1.**
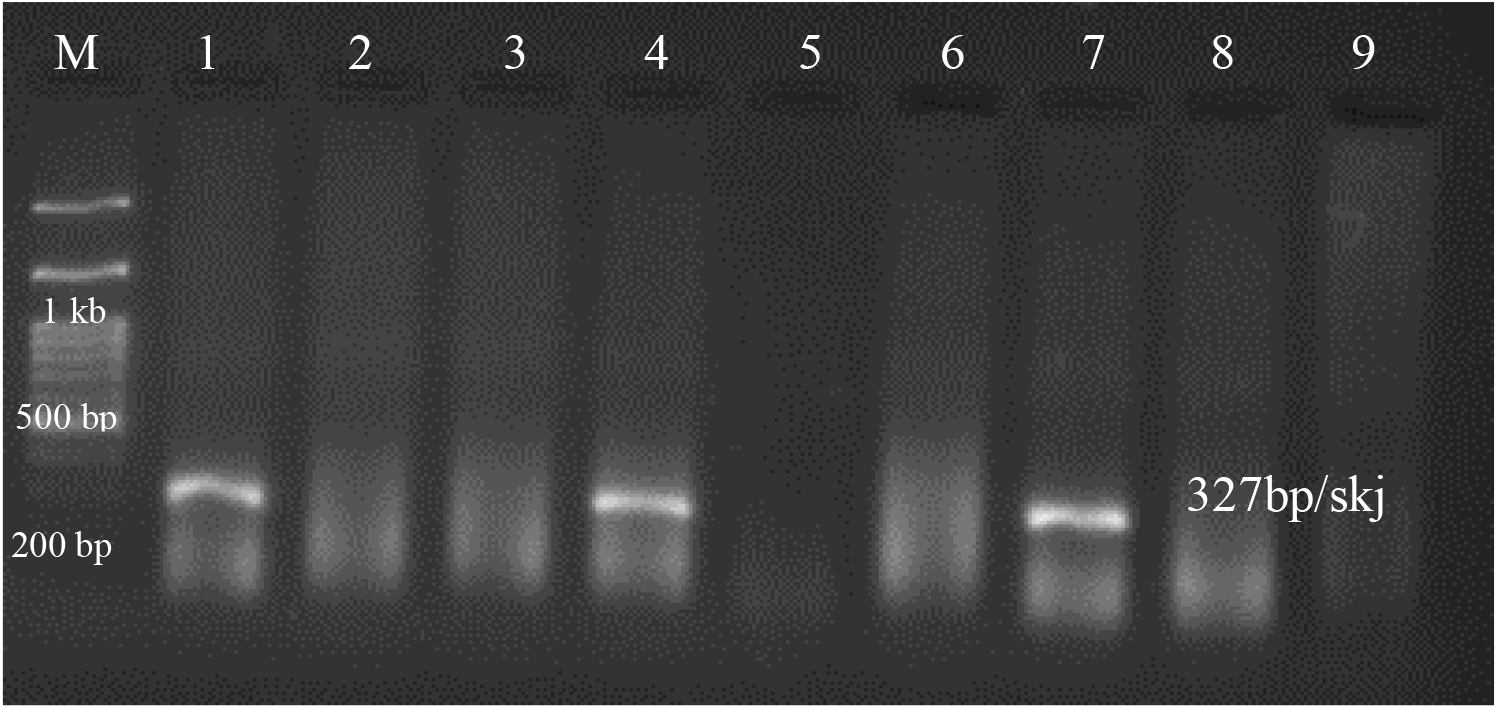
Las bandas de 327 bp por RT-PCR convencional *in house* del gen ORF7 (N). Línea 1 (ORF 7-SKJ): ADNc positivo cepa Nebraska Prime Pac^®^ PRRS-2; Línea 4 (ORF 7-SKJ): ADNc positiva cepa VP-046 SUIPRABAC^®^ PRRS-1; Línea 7 (ORF 7-SKJ): ADNc mezcla control positivo *Betaarterivirus americense* kit comercial IDEXX; Línea 2, 5 y 8 (bronquitis aviar): controles negativos de ADNc de IBV; Línea 3, 6 y 9 (blanco de PCR): agua libre de nucleasas; Línea M (ladder): 100 bp Plus Opti-DNA Marker ABM^®^.

### Phylogenetic Analysis of the Nucleocapsid (N) Protein

Phylogenetic relationships were inferred based on amino acid sequence alignment of the N protein, including representative GenBank strains from lineages 1, 5, and 8, together with the ten strains previously reported by Cotaquispe [5] (Figure 2). The maximum likelihood tree showed strong statistical support (100% bootstrap) for the separation between Betaarterivirus europensis and Betaarterivirus americense. Within PRRSV-2, distinct clades corresponding to lineage 1 (sublineages 1A [NADC34-like], 1C [NADC31-like], and 1C [HENAN-like]), lineage 5A (RespPRRS_MLV-like), and lineage 8 (Ingelvac ATP; JXA1-like) were identified. Eight strains from this study clustered within sublineage 1A (NADC34-like) (blue), whereas two strains grouped within lineage 5A (red) (Figure 2), demonstrating co-circulation of genetically distinct PRRSV-2 variants in the study region.

**Figure 2.**
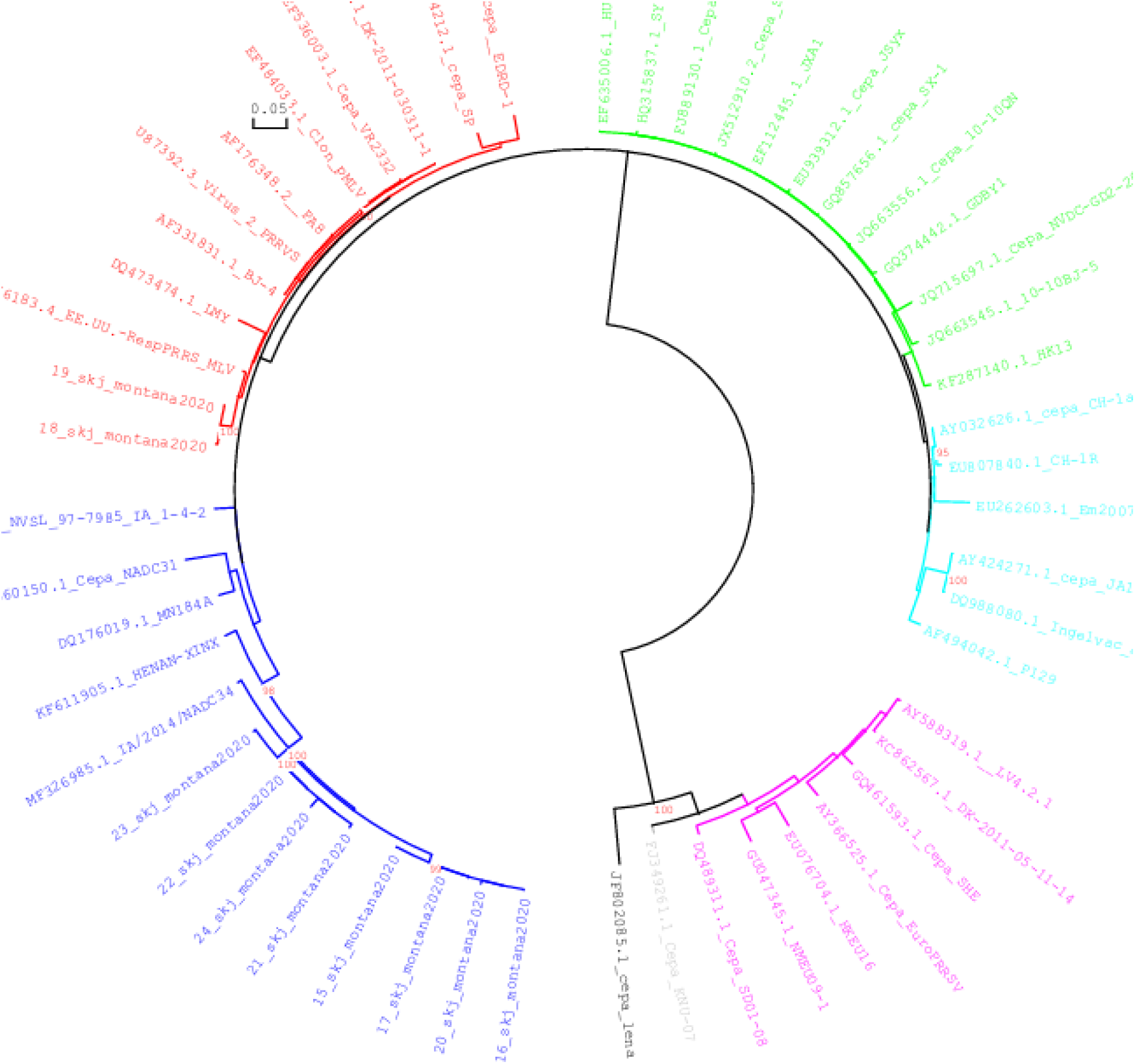
Árbol filogenético de las especies *Betaarterivirus europensis* (PRRSV-1) y *Betaarterivirus americense (*PRRSV-2) basado en la secuencia de la nucleocapside (N). (a) 1. Ramificación morada: 3 subtipos (*Betaarterivirus europensis*); 2. Ramificación azul (Linaje 1 *Betaarterivirus americense*): sublinaje 1A (NADC34 y 8 cepas estudio), sublinaje 1C (NADC31), linaje 1 (HENAN), linaje 1 (MN184A); 3. Ramificación roja (Linaje 5): L5A (Res-PRRS MLV, cepas 18-19 y VR2332); 4. Ramificación verde claro (Linaje 8): JXA1. (b) El análisis de sustitución de aminoácidos se realizó mediante el método de máxima verosimilitud aplicando BioNJ con el modelo Jones-Taylor-Thornton (JTT). Se utilizó el método bootstrap con 1000 réplicas de alineamiento y se realizaron análisis evolutivos en MEGA6 y DNAMAN 10.

**Figure 3.**
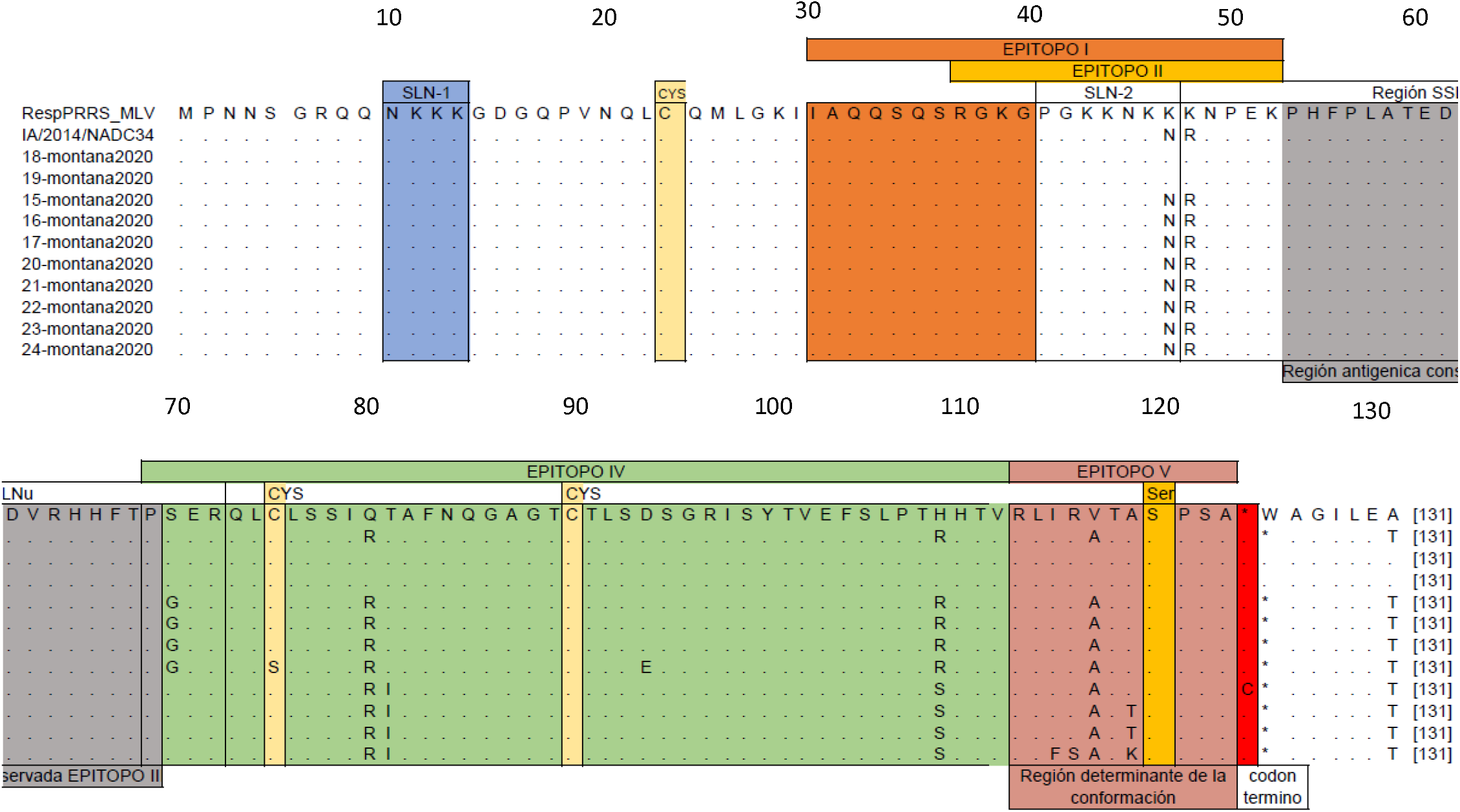
Alineamiento de las 10 secuencias (aa123) del gen ORF7 que codifica las regiones antigénicas de la nucleocápside (N) de *Betaarterivirus americense*. Las regiones enmarcadas y las coloreadas son 5 dominios inmunogénicos importantes pertenecientes a los linajes 1A y 5A identificados. Los 10 aislados de campo, las variantes A/2014/NADC34 (Linaje 1: sublinaje 1A) y RespPRRS_MLV (Linaje 5A) como patrones de referencia en el alineamiento local, los puntos indican la posición del aminoácido homologo, las letras distintas del punto en las cepas aisladas indican SNPs o cambios del aminoácido en comparación a las cepas de referencias.

### Genetic Diversity of the ORF7 (N) Gene

Amino acid sequences were aligned and compared with reference strains RespPRRS_MLV (lineage 5A) and A/2014/NADC34 (lineage 1A) to assess genetic variability. Strains 18 and 19 (lineage 5A) showed 100% amino acid identity with RespPRRS_MLV. In contrast, the eight lineage 1A strains exhibited multiple amino acid substitutions relative to A/2014/NADC34, particularly within antigenic domains IV and V. Within antigenic domain IV, substitutions included: S70G (4/8 strains), C75S (1/8), T81I (4/8), D94E (1/8), R109S (4/8); Within antigenic domain V: I115F (1/8), R116S (1/8), A119T/K (3/8). Strain 24 exhibited the highest divergence, accumulating five amino acid substitutions compared with the NADC34 reference strain. Despite this variability, conserved regions were observed within NLS-1 and the conserved cysteine (Cys23) region, indicating structural constraints within the N protein.

### Prediction of Linear B-Cell Epitopes

All ten strains were analyzed based on the 123-amino-acid ORF7 sequence to predict potential linear B-cell epitopes within the nucleocapsid protein. Six epitope patterns (A–F) were identified, comprising nine predicted B-cell epitope regions (positions 5–19, 33–72, 33–73, 84– 85, 87–98, 84–98, 87–97, 84, and 119). Pattern A corresponded exclusively to lineage 5A strains, whereas patterns B–F were associated with lineage 1A variants. Patterns B, E, and F displayed four to five predicted B-cell epitope regions and were observed in strains 21, 22, 23, and 24 (Table 1). These variations suggest potential modulation of antigenic profiles within lineage 1A variants.

**Table 1.**
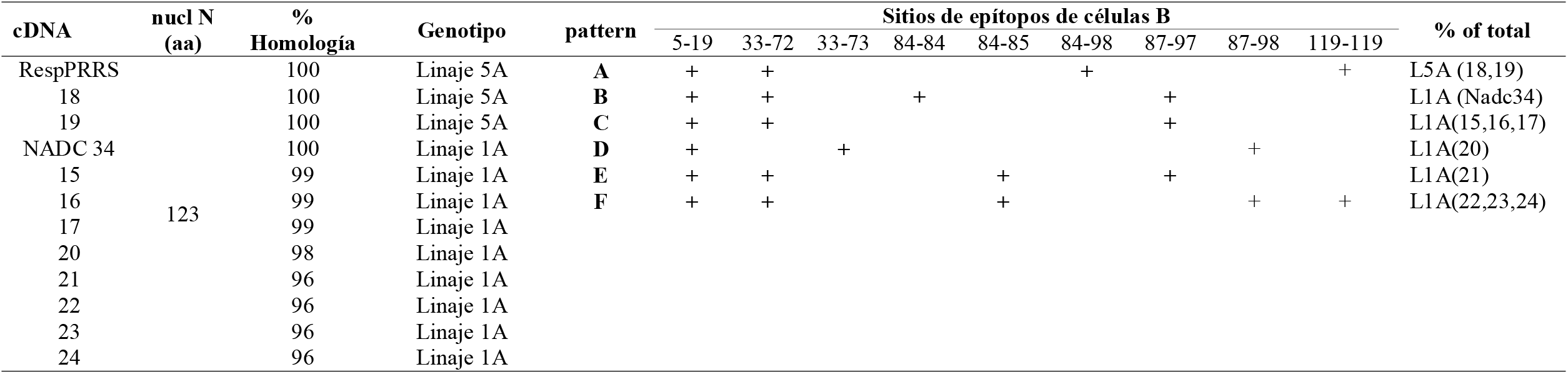
Análisis de variabilidad genotípica en las 10 secuencias de *Betaarterivirus americense* en la región de Lima, Perú.

### Genetic Diversity and Recombination Analysis

Genetic diversity analysis using DnaSP v6 identified 50 polymorphic nucleotide sites and 52 mutations within the ORF7 dataset (Table 2), indicating substantial sequence variability among the analyzed strains. Recombination analysis using RDP v4.101 detected a significant recombination event in strain 18_montana2020/L1A-R (highlighted in red; Figure 4). Breakpoints were located between nucleotide positions 141 and 414 of the alignment. The putative major parent was strain 24_montana2020/L1A-WT (green), whereas EF536003.1_L5A (vaccine-like VR2332 strain) was identified as the minor parent (blue) (Figure 4). This recombination event was supported by multiple independent detection methods (RDP, GENECONV, BootScan, MaxChi, Chimaera, SiScan, and 3Seq), with a highly significant Bonferroni-corrected global p-value of 1.441 × 10□^12^, minimizing the likelihood of false-positive detection. Phylogenetic trees constructed from non-recombinant regions (nt 1–140 and 415–523) and the central recombinant region (nt 141–414) showed a consistent topological shift for strain 18_montana2020. In flanking regions, the strain clustered with 24_montana2020, whereas in the recombinant central region it grouped with the lineage represented by EF536003.1_L5A (Table 3), confirming mosaic genome structure. Additionally, strain 19_montana2020/L5A-R exhibited high sequence identity with the recombinant strain and showed secondary evidence of the same recombination signal. However, it was not identified as the primary recombinant, suggesting either partial inheritance of the recombinant fragment or a shared ancestral recombination event (Figure 4).

**Table 2.**
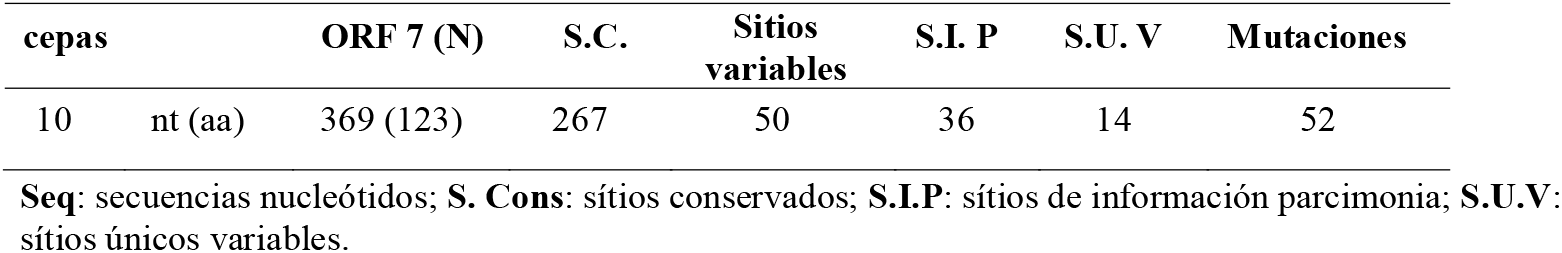
Análisis de diversidad genética de 10 aislados de *Betaarterivirus americense* analizados usando DnaSP y Mega 6 en muestras de sangre de cerdos de la región de Lima, Perú.

**Table 3.**
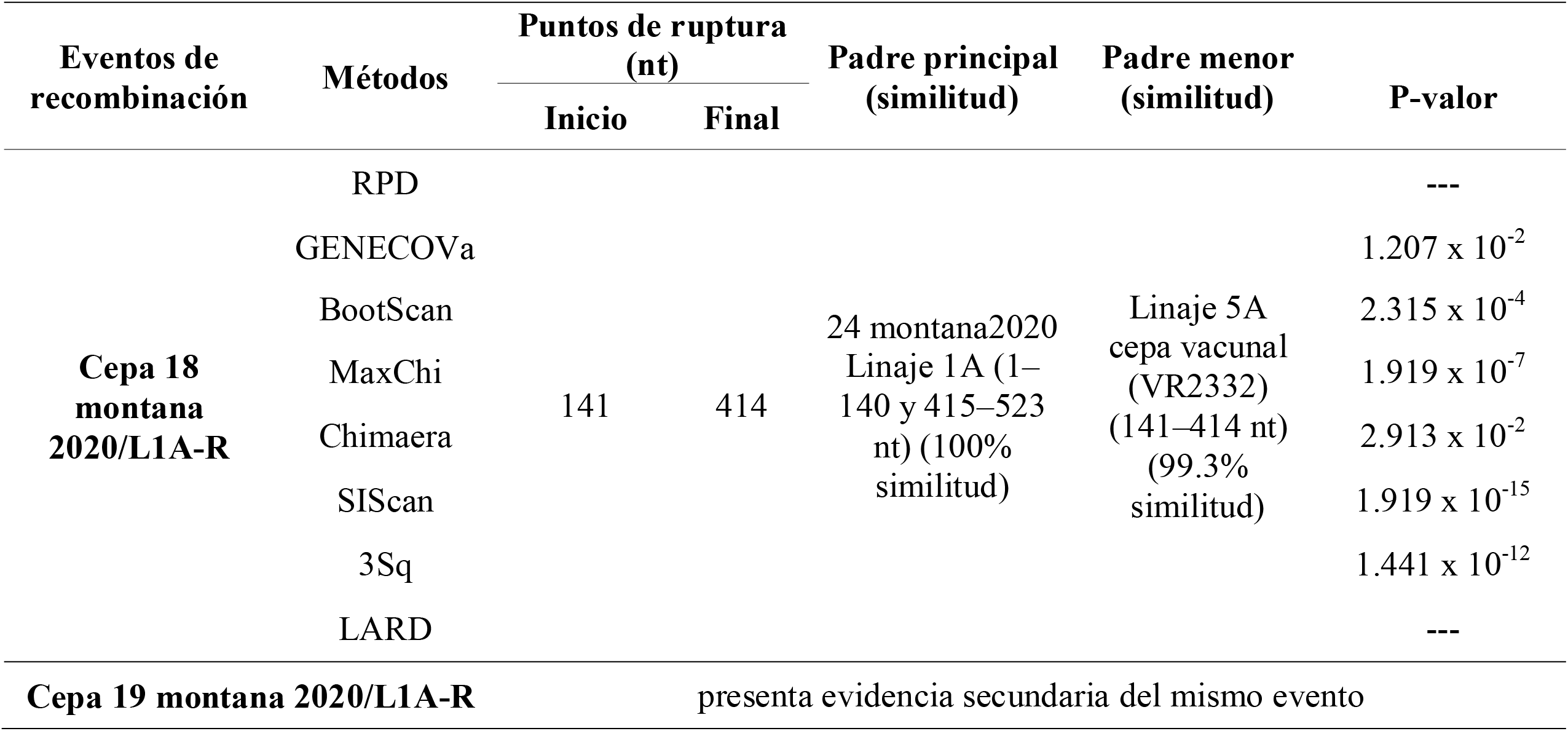
Genetic recombination events of Cepa 18 montana 2020-R and Cepa 19 montana 2020-R detected by software RDP versión 4.101.

**Figure 4.**
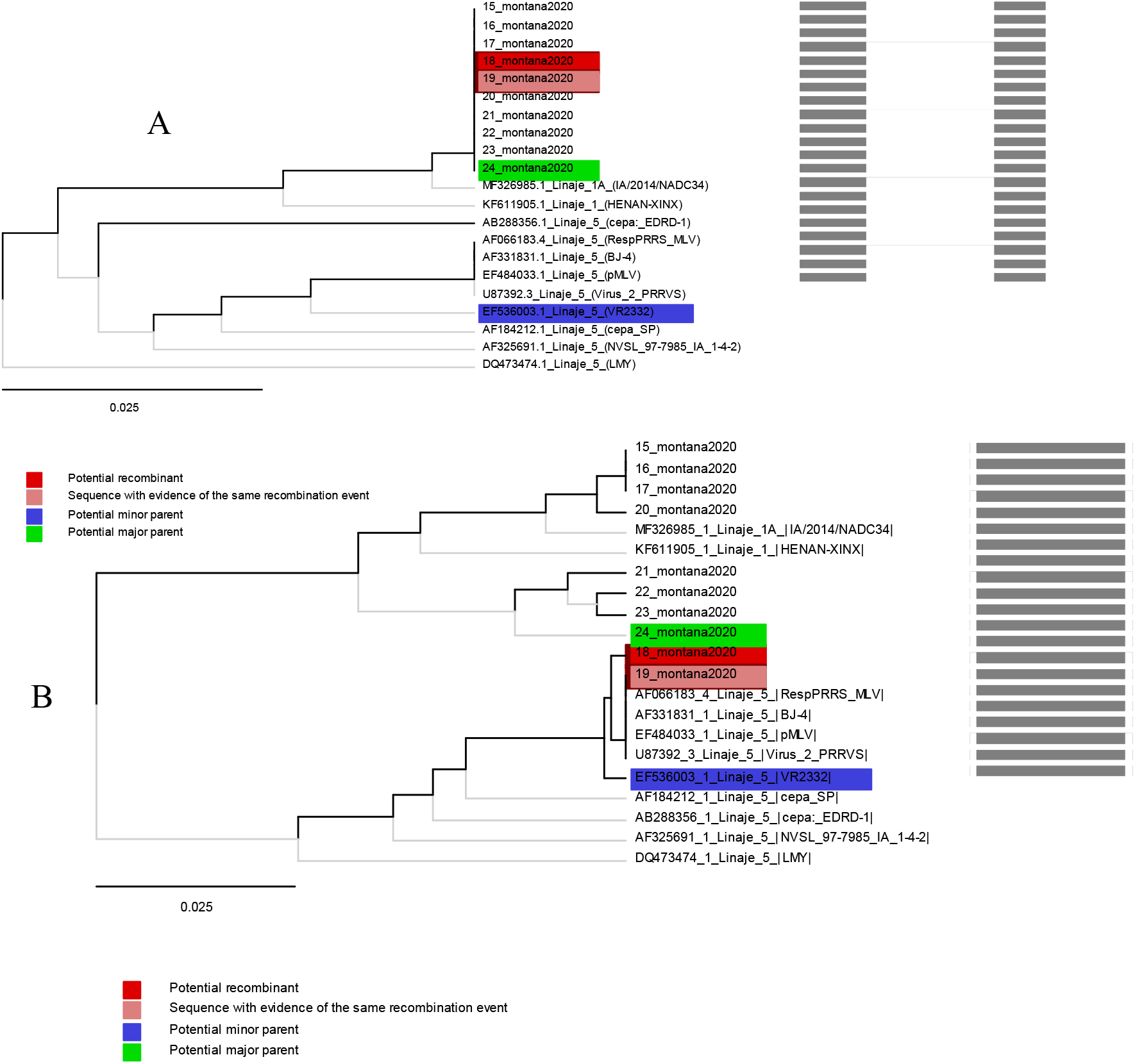
Análisis de recombinación genómica de la cepa 18 montana 2020-R (L1A) del virus del síndrome reproductivo y respiratorio porcino, Lima-Perú, 2026. A) UPGMA de la región derivada del progenitor principal 24 montana 2020/L1A (1–140 y 415–523 nt). B) UPGMA de la región derivada del progenitor menor Linaje 5A VR2332 (141–414 nt). Las filogenias de las cepas progenitoras se identificaron mediante el software RDP versión 4.101 (http://web.cbio.uct.ac.za/~darren/rdp.html). El rojo indica la cepa recombinante (18 montana 2020-R/L1A); el verde indica la cepa progenitora principal (cepa 24 montana 2020/L1A-WT); el azul indica la cepa progenitora secundaria Linaje 5A VR2332 (cepa vacunal). Las barras de escala indican las sustituciones de nucleótidos por sitio.

## DISCUSSION

The molecular analysis using the SKJ primers designed for the ORF7 gene proved highly effective for the detection and differentiation of Betaarterivirus europensis and Betaarterivirus americense lineages, successfully amplifying a 327 bp fragment corresponding to 369 nucleotides and encoding the full-length 123-amino-acid N protein. Among the ten sequences obtained, two belonged to lineage 5A, showing complete genetic identity with the RespPRRS_MLV vaccine strain, while the remaining eight clustered within lineage 1A (NADC34-like) [12, 35]. Notably, in a previous study, Cotaquispe [3] reported for the first time the presence of lineage 5A in pig farms in Lima, alongside substantial divergence among the eight 1A isolates. These observations underscore the need for further investigation into the genetic diversity and recombination potential within these circulating lineages.

Comparative alignment of ORF7 amino acid sequences revealed distinct substitution patterns among strains 15, 16, 17, 20, 21, 22, 23, and 24 relative to reference sequences. The distribution of substitutions is consistent with intra-lineage microevolution primarily driven by purifying selection, allowing the accumulation of structurally tolerable point mutations and the formation of discrete evolutionary subclusters. Strains 15–17 shared the S70G substitution, suggesting clonal expansion from a recent common ancestor. This change is compatible with neutral or nearly neutral evolution in structural proteins, particularly in regions that do not directly participate in critical oligomerization or RNA-binding interfaces. The conservation of S70G across multiple isolates indicates low functional constraint at this site.

Strain 20 exhibited an intermediate evolutionary profile, retaining S70G alongside additional substitutions C75S and D94E. While C75S may influence structural stability due to the role of cysteine residues in conformational integrity or intermolecular interactions, viral viability indicates that this position is not essential for nucleocapsid architecture. The D94E substitution represents a conservative change between negatively charged residues, consistent with genetic drift under functional constraints [8, 15, 21, 36, 37].

Strains 21–24 displayed a clearly differentiated pattern characterized by co-occurring T81I and R109S substitutions, suggesting fixation of a haplotype within a specific evolutionary subcluster. T81I involves a polar-to-hydrophobic transition, whereas R109S entails loss of positive charge; both could modulate local protein–protein or protein–RNA interactions without compromising overall stability. Additional substitutions, including A119T in strains 22 and 23 and I115F, R116S, and A119K in strain 24, indicate further subcluster-specific evolution. The clustering of mutations within the C-terminal region suggests relative structural flexibility or fine-tuning related to oligomerization or ribonucleoprotein assembly [28, 34, 37]. From a population perspective, shared mutational signatures reflect clonal expansion dynamics, local transmission, and potential population bottlenecks. This pattern is typical of essential structural viral genes, where functional conservation restricts disruptive mutations while allowing evolutionary plasticity. Overall, these results validate ORF7 as a reliable molecular marker for short-term epidemiological studies, enabling sublineage discrimination while preserving robust phylogenetic comparability.

Predicted linear B-cell epitope analysis demonstrated evolutionary patterns consistent with highly conserved structural viral proteins, where stable immunodominant regions coexist with antigenically variable zones. Six epitope patterns (A–F) with nine predicted B-cell epitope sites (positions 5–19, 33–72, 33–73, 84–85, 87–98, 84–98, 87–97, 84, and 119) suggest strong purifying selection, likely reflecting structural and functional importance within the N protein architecture [2]. In contrast, patterns B, E, and F, found in strains 21–24, indicate regions with higher evolutionary tolerance. This observation is consistent with localized antigenic diversification, whereby the virus maintains functionally constrained epitopes while generating variability in peripheral regions.

The increased number of potential epitopes in strains 21–24 further supports intra-lineage microdiversification, potentially driven by differential immune pressure favoring variants with slightly altered antigenic profiles. Alternatively, this may represent an antigenic redistribution strategy, whereby new immunogenic determinants divert humoral responses away from functionally critical regions [30]. Collectively, these findings suggest that the N protein maintains essential structural roles while indirectly influencing virus–host interactions through modulation of its antigenic landscape, reinforcing the importance of considering N protein variability in viral evolution studies and serological interpretations.

Our results indicate that ORF7 genetic diversity among the Montana 2020 isolates is generally low, consistent with recent circulation or local transmission within a single lineage. The monophyletic clustering and short genetic distances observed among sequences 15– 24_montana2020 imply a common origin and limited accumulation of point mutations. Despite this apparent homogeneity, recombination analysis using RDP v4.101 identified a statistically robust homologous recombination event in strain 18_montana2020/L5A-R, with well-defined breakpoints (nt 141–414) and strong support across multiple independent algorithms, including RDP, GENECONV, Bootscan, MaxChi, Chimaera, SiScan, and 3Seq (Bonferroni-corrected p = 1.441 × 10□^12^) [12, 19, 32].

Phylogenetic comparison of recombinant and non-recombinant regions demonstrated a clear mosaic pattern: strain 18_montana2020/L5A-R clustered with the Montana 2020 clade in flanking regions but aligned with the EF536003_1_L5A vaccine-like lineage in the central ORF7 region. This independent phylogenetic evidence confirms the recombinant origin. Identification of strain 24_montana2020/L1A-WT as the putative major parent and a divergent lineage 5A strain as the minor parent indicates co-circulation of genetically distinct variants and possible coinfections that facilitated genetic exchange. Notably, strain 19_montana2020/L5A-R exhibited secondary evidence of the same recombination event, consistent with clonal propagation of an already recombined genome rather than independent recombination events. This finding has epidemiological implications, demonstrating that recombination can not only occur but also persist and disseminate in local viral populations [12, 23, 32].

From an evolutionary perspective, detection of recombination within ORF7 is particularly relevant due to the functional role of the N protein in viral assembly, genome encapsidation, and host immune modulation. Although ORF7 is generally considered relatively conserved, our results demonstrate that it can serve as a substrate for homologous recombination, generating mosaic variants with potential adaptive advantages. Overall, these findings highlight that even in contexts of apparently low genetic diversity, recombination can play a critical role in the evolutionary dynamics of Betaarterivirus americense, increasing genetic variability and adaptive potential. Consequently, molecular surveillance and phylodynamic reconstruction studies should systematically incorporate recombination analyses to avoid biased phylogenetic inferences and more accurately understand the mechanisms driving viral evolution.

## CONCLUSIONS

✓ Phylogenetic analysis using DNAMAN v10.0 and MEGA 6 identified *Betaarterivirus americense* in the ten isolates, revealing the presence of two lineages: eight strains clustered within lineage 1A (NADC34) and two strains within lineage 5A (VR2322), based on ORF7 nucleocapsid (N) gene alignment.
✓ Comparative amino acid alignment in MEGA 6 revealed significant substitutions in strains 15, 16, 17, 20, 21, 22, 23, and 24, with strain 24 being the most divergent. Multiple C-terminal substitutions were observed (T81I, R109S, I115F, R116S, and A119K), located within the N protein antigenic domains I–V.
✓ B-cell epitope prediction using BepiPred-2.0 identified six patterns (A–F) with nine potential linear B-cell epitope sites (positions 5–19, 33–72, 33–73, 84–85, 87–98, 84– 98, 87–97, 84, and 119), with patterns B, E, and F containing four to five epitope sites corresponding to strains 21, 22, 23, and 24.
✓ Recombination analysis using RDP v4.101 detected a statistically robust homologous recombination event in strain 18_montana2020-R (lineage 5A), with the major parent identified as strain 24_montana2020-WT (lineage 1A, 100% similarity) and the minor parent as EF536003_1_L5A (vaccine strain VR2322, 99.3% similarity). Strain 19_montana2020-R showed secondary evidence of the same recombination event.
✓ Genetic diversity analysis of the ORF7 (N) sequences revealed 50 polymorphic nucleotide sites and 52 mutations, indicating significant intra-lineage variability.

## ACKNOWLEDGMENTS

The authors sincerely thank Corporación Montana S.A. for funding and providing logistical support for this research. We also acknowledge Bach. Segundo Del Águila Soto for his valuable technical assistance and insightful suggestions.

