## Supplementary material for "Molecular Diversity and Recombination Patterns of the ORF7 (Nucleocapsid) Gene in *Betaarterivirus americense* Variants Circulating from Lima, Peru": Material suplementary

>15_montana2020

AATATGCCAAATAACAGCGGCAGACAGCAAAATAAAAAGAAGGGGGATGGCCAGCCAGTCAATCAGCTGT
GCCAGATGTTGGGCAAGATTATCGCCCAACAGAGCCAGTCCAGAGGCAAGGGACCGGGGAAGAAGAACA
AGAATAGGAACCCGGAGAAGCCCCATTTTCCTCTAGCGACTGAAGATGACGTCAGACATCACTTTACCCCCG
GTGAGCGACAATTGTGTCTGTCGTCAATCCGGACTGCCTTTAACCAAGGCGCTGGAACTTGTACCCTGTCAG
ATTCAGGGAGAATAAGTTACACTGTGGAATTCAGTCTGCCTACTCGTCATACTGTGCGCTTGATTCGCGCCA
CAGCGTCACCCTCAGCATGATGAGCTGGCATTCTTGAGACATCCCAGTGCCTGAATTGGAAGAATGAGTGG
TGAACGGCACTGATTGACATTGTGCCTCTAAGTCACCTATTCAATTAGGGCGACCGTATGGGGGTAATATTT
AATTGGCGAGAACCATGCGGCCGAA

>16_montana2020

AATATGCCAAATAACAGCGGCAGACAGCAAAATAAAAAGAAGGGGGATGGCCAGCCAGTCAATCAGCTGT
GCCAGATGTTGGGCAAGATTATCGCCCAACAGAGCCAGTCCAGAGGCAAGGGACCGGGGAAGAAGAACA
AGAATAGGAACCCGGAGAAGCCCCATTTTCCTCTAGCGACTGAAGATGACGTCAGACATCACTTTACCCCCG
GTGAGCGACAATTGTGTCTGTCGTCAATCCGGACTGCCTTTAACCAAGGCGCTGGAACTTGTACCCTGTCAG
ATTCAGGGAGAATAAGTTACACTGTGGAATTCAGTCTGCCTACTCGTCATACTGTGCGCTTGATTCGCGCCA
CAGCGTCACCCTCAGCATGATGAGCTGGCATTCTTGAGACATCCCAGTGCCTGAATTGGAAGAATGAGTGG
TGAACGGCACTGATTGACATTGTGCCTCTAAGTCACCTATTCAATTAGGGCGACCGTATGGGGGTAATATTT
AATTGGCGAGAACCATGCGGCCGAA

>17_montana2020

AATATGCCAAATAACAGCGGCAGACAGCAAAATAAAAAGAAGGGGGATGGCCAGCCAGTCAATCAGCTGT
GCCAGATGTTGGGCAAGATTATCGCCCAACAGAGCCAGTCCAGAGGCAAGGGACCGGGGAAGAAGAACA
AGAATAGGAACCCGGAGAAGCCCCATTTTCCTCTAGCGACTGAAGATGACGTCAGACATCACTTTACCCCCG
GTGAGCGACAATTGTGTCTGTCGTCAATCCGGACTGCCTTTAACCAAGGCGCTGGAACTTGTACCCTGTCAG
ATTCAGGGAGAATAAGTTACACTGTGGAATTCAGTCTGCCTACTCGTCATACTGTGCGCTTGATTCGCGCCA
CAGCGTCACCCTCAGCATGATGAGCTGGCATTCTTGAGACATCCCAGTGCCTGAATTGGAAGAATGAGTGG
TGAACGGCACTGATTGACATTGTGCCTCTAAGTCACCTATTCAATTAGGGCGACCGTATGGGGGTAATATTT
AATTGGCGAGAACCATGCGGCCGAA

>18_montana2020

AATATGCCAAATAACAGCGGCAGACAGCAAAATAAAAAGAAGGGGGATGGCCAGCCAGTCAATCAGCTGT
GCCAGATGTTGGGCAAGATTATCGCCCAACAGAGCCAGTCCAGAGGCAAGGGACCGGGGAAGAAGAACA
AGAAGAAAAACCCGGAGAAGCCCCATTTTCCTCTAGCGACTGAAGATGATGTCAGACATCATTTTACCCCTA
GTGAGCGGCAATTGTGTCTGTCGTCAATCCAGACCGCCTTTAATCAAGGCGCTGGGACTTGCACCCTGTCAG
ATTCAGGGAGGATAAGTTACACTGTGGAGTTTAGTTTGCCTACGCATCATACTGTGCGCCTGATCCGCGTCA
CAGCATCACCCTCAGCATGATGGGCTGGCATTCTTGAGGCATCTCAGTGTTTGAATTGGAAGAATGAGTGG
TGAACGGCACTGATTGACATTGTGCCTCTAAGTCACCTATTCAATTAGGGCGACCGTATGGGGGTAATATTT
AATTGGCGAGAACCATGCGGCCGAA

>19_montana2020

AATATGCCAAATAACAGCGGCAGACAGCAAAATAAAAAGAAGGGGGATGGCCAGCCAGTCAATCAGCTGT
GCCAGATGTTGGGCAAGATTATCGCCCAACAGAGCCAGTCCAGAGGCAAGGGACCGGGGAAGAAGAACA

AGAAGAAAAACCCGGAGAAGCCCCATTTTCCTCTAGCGACTGAAGATGATGTCAGACATCACTTTACCCCTA
GTGAGCGGCAATTGTGTCTGTCGTCAATCCAGACCGCCTTTAATCAAGGCGCTGGGACTTGCACCCTGTCAG
ATTCAGGGAGGATAAGTTACACTGTGGAGTTTAGTTTGCCTACGCATCATACTGTGCGCCTGATCCGCGTCA
CAGCATCACCCTCAGCATGATGGGCTGGCATTCTTGAGGCATCTCAGTGTTTGAATTGGAAGAATGAGTGG
TGAACGGCACTGATTGACATTGTGCCTCTAAGTCACCTATTCAATTAGGGCGACCGTATGGGGGTAATATTT
AATTGGCGAGAACCATGCGGCCGAA

>20_montana2020

AATATGCCAAATAACAGCGGCAGACAGCAAAATAAAAAGAAGGGGGATGGCCAGCCAGTCAATCAGCTGT
GCCAGATGTTGGGCAAGATTATCGCCCAACAGAGCCAGTCCAGAGGCAAGGGACCGGGGAAGAAGAACA
AGAATAGGAACCCGGAGAAGCCCCATTTTCCTCTAGCGACTGAAGATGACGTCAGACATCACTTTACCCCCG
GTGAGCGACAATTGAGTCTGTCGTCAATCCGGACTGCCTTTAACCAAGGCGCTGGAACTTGTACCCTGTCAG
AATCAGGGAGAATAAGTTACACTGTGGAATTCAGTCTGCCTACTCGTCATACTGTGCGCTTGATTCGCGCCA
CAGCGTCACCCTCAGCATGATGAGCTGGCATTCTTGAGACATCCCAGTGCCTGAATTGGAAGAATGAGTGG
TGAACGGCACTGATTGACATTGTGCCTCTAAGTCACCTATTCAATTAGGGCGACCGTATGGGGGTAATATTT
AATTGGCGAGAACCATGCGGCCGAA

>21_montana2020

AATATGCCAAATAACAGCGGCAGACAGCAAAATAAAAAGAAGGGGGATGGCCAGCCAGTCAATCAGCTGT
GCCAGATGTTGGGCAAGATTATCGCCCAACAGAGCCAGTCCAGAGGCAAGGGACCGGGGAAGAAGAACA
AGAACAGAAACCCGGAGAAGCCCCATTTTCCTCTAGCGACTGAAGATGACGTCAGACATCACTTTACCCCCA
GTGAACGACAATTGTGTCTGTCGTCAATCCGAATTGCCTTTAACCAAGGCGCTGGAACTTGTACCCTGTCAG
ACTCGGGGAGAATAAGTTACACTGTGGAGTTCAGTCTGCCTACTAGTCATACTGTGCGTTTGATTCGCGCCA
CAGCGTCACCATCAGCATGCTGAGCTGGCATTCTTGAGACATCCCAGTGCTTGAATTGGAAGAATGAGTGG
TGAACGGCACTGATTGACATTGTGCCTCTAAGTCACCTATTCAATTAGGGCGACCGTATGGGGGTAATATTT
AATTGGCGAGAACCATGCGGCCGAA

>22_montana2020

AATATGCCAAATAACAGCGGCAGACAGCAAAATAAAAAGAAGGGGGATGGCCAGCCAGTCAATCAGCTGT
GCCAGATGTTGGGCAAGATTATCGCCCAACAGAGCCAGTCCAGAGGCAAGGGACCGGGGAAGAAGAACA
AGAACAGAAACCCGGAGAAGCCCCATTTTCCTCTAGCGACTGAAGATGACGTCAGACATCACTTTACCCCCA
GTGAACGACAATTGTGTCTGTCGTCAATCCGAATTGCCTTTAACCAAGGCGCTGGAACTTGTACCCTGTCAG
ACTCGGGGAGAATAAGTTACACTGTGGAGTTCAGTCTGCCTACTAGTCATACTGTGCGTTTGATTCGCGCCA
CAACGTCACCCTCAGCATGATGAGCTGGCATTCTTGAGACATCCCAGTGCTTGAATTGGAAGAATGAGTGG
TGAACGGCACTGATTGACATTGTGCCTCTAAGTCACCTATTCAATTAGGGCGACCGTATGGGGGTAATATTT
AATTGGCGAGAACCATGCGGCCGAA

>23_montana2020

AATATGCCAAATAACAGCGGCAGACAGCAAAATAAAAAGAAGGGGGATGGCCAGCCAGTCAATCAGCTGT
GCCAGATGTTGGGCAAGATTATCGCCCAACAGAGCCAGTCCAGAGGCAAGGGACCGGGGAAGAAGAACA
AGAACAGAAACCCGGAGAAGCCCCATTTTCCTCTAGCGACTGAAGATGACGTCAGACATCACTTTACCCCCA
GTGAACGACAATTGTGTCTGTCGTCAATCCGGATTGCCTTTAACCAAGGCGCTGGAACTTGTACCCTGTCAG
ACTCGGGGAGAATAAGTTACACTGTGGAGTTCAGTCTGCCTACTAGTCATACTGTGCGTTTGATTCGCGCCA
CAACGTCACCCTCAGCATGATGAGCTGGCATACTTGAGACATCCCAGTGCTTGAATTGGAAGAATGAGTGG

TGAACGGCACTGATTGACATTGTGCCTCTAAGTCACCTATTCAATTAGGGCGACCGTATGGGGGTAATATTT
AATTGGCGAGAACCATGCGGCCGAA

>24_montana2020

AATATGCCAAATAACAGCGGCAGACAGCAAAATAAAAAGAAGGGGGATGGCCAGCCAGTCAATCAGCTGT
GCCAGATGTTGGGCAAGATTATCGCCCAACAGAGCCAGTCCAGAGGCAAGGGACCGGGGAAGAAGAACA
AGAACAGAAACCCGGAGAAGCCCCATTTTCCTCTAGCGACTGAAGATGACGTCAGACATCACTTTACCCCCA
GTGAACGACAATTGTGTCTGTCGTCAATCCGGATTGCCTTCAACCAAGGCGCTGGAACTTGTACCCTGTCAG
ACTCGGGGAGAATAAGTTACACTGTGGAGTTCAGTCTGCCTACTAGTCATACTGTGCGTTTGTTTAGCGCCA
CGAAGTCACCATCAGCATGATGAGCTGGCATTCTTGAGACATCCCAGTGCTTGAATTGGAAGAATGAGTGG
TGAACGGCACTGATTGACATTGTGCCTCTAAGTCACCTATTCAATTAGGGCGACCGTATGGGGGTAATATTT
AATTGGCGAGAACCATGCGGCCGAA

>MF326985.1_Linaje_1A_(IA/2014/NADC34)

AATATGCCAAATAACAGCGGCAGACAGCAAAATAAAAAGAAGGGGGATGGCCAGCCAGTCAATCAGCTGT
GCCAGATGTTGGGCAAGATTATCGCCCAACAGAGCCAGTCCAGAGGCAAGGGACCGGGGAAGAAGAATA
AGAATAGAAACCCGGAGAAGCCCCATTTTCCTCTAGCGACTGAAGATGACGTCAGACATCACTTTACCCCCA
GTGAGCGACAATTGTGTCTGTCGTCAATCCGGACTGCCTTTAACCAAGGCGCTGGAACTTGCACCCTGTCAG
ACTCAGGGAGAATAAGTTACACTGTGGAGTTCAGTCTGCCTACTCGTCATACTGTGCGCTTGATTCGCGCCA
CAGCGTCACCCTCAGCATGATGAGCTGGCATTCTTGAGACATCCCAGTGCCTGAATTGGAAGAATGAGTGG
TGAATGGCACTGATTGACATTGTGCCTCTAAGTCACCTATTCAATTAGGGCGACCGTATGGGGGTAATATTT
AATTGGCGAGAACCATGCGGCCGAA

>KF611905.1_Linaje_1_(HENAN-XINX)

AATATGCCAAATAACAACGGCAGACAGCAAAATAAAAAGAAGGGGGATGGCCAGCCAGTCAATCAGCTGT
GCCAGATGCTGGGTAAGATTATTGCCCAACAGAGCCAGTCCAGAGGCAAGGGACCGGGAAAGAAAAATAA
GAATAGAAGCCCGGAGAAGCCCCATTTTCCTCTAGCGACTGAAGATGACGTCAGACATCACTTTACCCCTAG
TGAGCGACAATTGTGTCTGTCGTCAATCCGGACTGCCTTTAACCAAGGCGCTGGAACTTGTACCCTGTCAGA
CTCAGGGAGAATAAGTTACACTGTGGAGTTTAGTTTGCCTACTCATCATACCGTGCGCCTGATTCGCGCCAC
AGCGTCACCCTCAGCATGATGAGCTGGCATTCTTGAAACATCCCGGTGTTTGAATTGGAAGAATGAGTGGT
GAATGGCACTGATTGATATTGTGCCTCTAAGTCACCTATTCAATTAGGGCGACCGTATGGGGGTAACATTTA
ATTGGCGAGAACCATGCGGCCGAA

>AB288356.1_Linaje_5_(cepa:_EDRD-1)

AATATGCCAAATAACAACGGCAAACAGCAAAAGAGAAAGAAGGGGAATGGCCAGCCAGTCAATCAGCTGT
GCCAGATGCTGGGTAAGATTATTGCCCAGCAGAGTCAGTCCAGAGGTAAGGGACCGGGAAATAAAAACAA
GAAGAAAAATCCGGAGAAGCCCCATTTTCCTCTAGCGACTGAATATGACGTCAGACATCACTTCACCCCTAG
TGAGCGGCAATTGTGTCTGTCGTCAATCCAGACTGCCTTTAATCAAGGCGCTGGAACTTGCACCCTGTCAGA
TTCAGGGAGGATAAGTTACACTGTGGAGTTTAGTTTGCCTACGCATCATACCGTGCGCCTTATTCGCGTCAC
AGCATCACCCTCAGCATGATGAGCTGGCATTCTTGAGGCACCTCAGTGTTTAAATTGGAAGAATGTGTGGT
GAATGGCACTGATTGATATTGTGCCTCTAAGTCACCTATTCAATTAGGGCGACCGTGTGGGGGTAGGATTT
AATTGGCGATAACCACGCGGCCGAA

>AF066183.4_Linaje_5_(RespPRRS_MLV)

AATATGCCAAATAACAACGGCAAGCAGCAGAAGAGAAAGAAGGGGGATGGCCAGCCAGTCAATCAGCTG
TGCCAGATGCTGGGTAAGATCATCGCTCAGCAAAACCAGTCCAGAGGCAAGGGACCGGGAAAGAAAAATA
AGAAGAAAAACCCGGAGAAGCCCCATTTTCCTCTAGCGACTGAAGATGATGTCAGACATCACTTTACCCCTA
GTGAGCGGCAATTGTGTCTGTCGTCAATCCAGACCGCCTTTAATCAAGGCGCTGGGACTTGCACCCTGTCAG
ATTCAGGGAGGATAAGTTACACTGTGGAGTTTAGTTTGCCTACGCATCATACTGTGCGCCTGATCCGCGTCA
CAGCATCACCCTCAGCATGATGGGCTGGCATTCTTGAGGCATCTCAGTGTTTGAATTGGAAGAATGTGTGG
TGAATGGCACTGATTGACATTGTGCCTCTAAGTCACCTATTCAATTAGGGCGACCGTGTGGGGGTGAGATTT
AATTGGCGAGAACCATGCGGCCGAA

>AF184212.1_Linaje_5_(cepa_SP)

AATATGCCAAATAACAACGGCAAACAGCAGAAGAAAAAGAAGGGGGATGGCCAGCCAGTCAATCAGCTGT
GCCAGATGCTGGGTAAGATCATCGCCCAGCAAAACCAGTCCAGAGGTAAGGGACCGGGAAAGAAAAACA
AGAAGAAAAACCCGGAGAAGCCCCATTTTCCTCTGGCGACTGAATATGACGTCAGACACCACTTTACCCCTA
GTGAGCGGCAATTGTGCCTGTCGTCAATACAGACTGCCTTTAATCAAGGCGCTGGTACTTGCACCCTGTCCG
ATTCAGGGAGGATAAGTTACACTGTGGAGTTTAGTTTGCCCACGCATCATACTGTGCGCCTGATTCGCGTCA
CAGCATCACCCTCAGCATGATGGGCTGGCATTCTTGAGGCATCTCAGTGTTTGAATTGGAAGAATGTGTGG
TGAATGGCACTGATTGATATTGTGCCTCTAAGTCACCTATTCAATTAGGGCGACCGTGTGGGGGTAAGATTT
AATTGGCGAAAACCATGCGGCCGAA

>AF325691.1_Linaje_5_(NVSL_97-7985_IA_1-4-2)

AATATGCCAAATAACAACGGCAAGCAGCAAAAGAAAAAGAAGGGGAATGGCCAGCCAGTCAACCAGCTGT
GCCAAATGCTGGGCAAGATCATCGCCCAGCAGAACCAGTCCAGAGGTAAGGGACCGGGAAAGAAAATTAA
AAAGAAAAACCCGGAGAAGCCCCATTTTCCTCTAGCGACCGAAGATGACGTCAGACATCACTTTACCCCTAG
TGAGCGGCAATTGTGTCTGTCGTCAATCCAGACTGCCTTTAATCAAGGCGCTGGAACTTGCACCCTGTCGGA
TTCAGGGAGGATAAGTTACACTGTGGAGTTTAGTTTGCCGACGCATCATACTGTGCGCCTGATTCGCGTCAC
AGCACCACCCTCAGCGTGATGGGCTGGCATTCTTGAGACATCCCAGTGTTAGAATTGGAAGAATGTGTGGT
GAATGGCACTGATTGACACTGTGCCTCTAAGTCACCTATTCAATTAGGGCGACCGTGTGGGGGTGAGATTT
AATTGGCGAGAACCATGCGGCCGAA

>AF331831.1_Linaje_5_(BJ-4)

AATATGCCAAATAACAACGGCAAGCAGCAGAAGAGAAAGAAGGGGGATGGCCAGCCAGTCAATCAGCTG
TGCCAGATGCTGGGTAAGATCATCGCTCAGCAAAACCAGTCCAGAGGCAAGGGACCGGGAAAGAAAAATA
AGAAGAAAAACCCGGAGAAGCCCCATTTTCCTCTAGCGACTGAAGATGATGTCAGACATCACTTTACCCCTA
GTGAGCGGCAATTGTGTCTGTCGTCAATCCAGACCGCCTTTAATCAAGGCGCTGGGACTTGCACCCTGTCAG
ATTCAGGGAGGATAAGTTACACTGTGGAGTTTAGTTTGCCTACGCATCATACTGTGCGCCTGATCCGCGTCA
CAGCATCACCCTCAGCATGATGGGCTGGCATTCTTGAGGCATCTCAGTGTTTGAATTGGAAGAATGTGTGG
TGAATGGCACTGATTGACATTGTGCCTCTAAGTCACCTATTCAATTAGGGCGACCGTGTGGGGGTGAGATTT
AATTGGCGAGAACCATGCGGCCGAA

>DQ473474.1_Linaje_5_(LMY)

AATATGCCAAATAACAACGGCAAGCAGCAGAAGAGAAAGAAGGGGGATGGCCAGCCAGTCAATCAGCTG
TGCCAAATGCTGGGCAAAATTATCGCCCAGCAAAACCAGTCCAGAGGCAAGGGACCGGGAAAGAAAAATA
AGAAGAAAAACCCCGAGAAGCCCCATTTCCCTCTTGCGGCTGAAGATGATGTCAGACATCACTTTACCCCTA
GTGAGCGGCAATTGTGTCTGTCGTCAATCCAGACCGCCTTTAATCAAGGCGCTGGGACCTGCACCCTATCAG

ATTCAGGGAGGATAAGTTACACTGTGGAGTTTAGTTTGCCTACGCATCATACTGTGCGCCTGATCCGTGTCA
CAGCATCACCTTCAGCATGATGAGCTGGCATTCTTGTGGCATCCCAGTATTTGAATTGGAATAATGCGTGGT
GAATGGCACTGATTGACATTGTGCCTTCAAGTCACCTATTCAATTAGGGCGACCGTGTGGGGGCAAGATTT
AATTGGCGAGAACCACACGGCCGAA

>EF484033.1_Linaje_5_(pMLV)

AATATGCCAAATAACAACGGCAAGCAGCAGAAGAGAAAGAAGGGGGATGGCCAGCCAGTCAATCAGCTG
TGCCAGATGCTGGGTAAGATCATCGCTCAGCAAAACCAGTCCAGAGGCAAGGGACCGGGAAAGAAAAATA
AGAAGAAAAACCCGGAGAAGCCCCATTTTCCTCTAGCGACTGAAGATGATGTCAGACATCACTTTACCCCTA
GTGAGCGGCAATTGTGTCTGTCGTCAATCCAGACCGCCTTTAATCAAGGCGCTGGGACTTGCACCCTGTCAG
ATTCAGGGAGGATAAGTTACACTGTGGAGTTTAGTTTGCCTACGCATCATACTGTGCGCCTGATCCGCGTCA
CAGCATCACCCTCAGCATGATGGGCTGGCATTCTTGAGGCATCTCAGTGTTTGAATTGGAAGAATGTGTGG
TGAATGGCACTGATTGACATTGTGCCTCTAAGTCACCTATTCAATTAGGGCGACCGTGTGGGGGTGAGATTT
AATTGGCGAGAACCATGCGGCCGAA

>EF536003.1_Linaje_5_(VR2332)

AATATGCCAAATAACAACGGCAAGCAGCAGAATAGAAAGAAGGGGGATGGCCAGCCAGTCAATCAGCTGT
GCCAGATGCTGGGTAAGATCATCGCTCAGCAAAACCAGTCCAGAGGCAAGGGACCGGGAAAGAAAAATAA
GAAGAAAAACCCGGAGAAGCCCCATTTTCCTCTAGCGACTGAAGATGATGTCAGACATCACTTTACCCCTAG
TGAGCGGCAATTGTGTCTGTCGTCAATCCAGACCGCCTTTAATCAAGGCGCTGGGACTTGCACCCTGTCAGA
TTCAGGGAGGATAAGTTACACTGTGGAGTTTAGTTTGCCTACGCATCATACTGTGCGCCTGATTCGCGTCAC
AGCATCACCCTCAGCATGATGGGCTGGCATTCTTGAGGCATCTCAGTGTTTGAATTGGAAGAATGTGTGGT
GAATGGCACTGATTGACATTGTGCCTCTAAGCACTATATT-----------------------------------------------------

>U87392.3_Linaje_5_(Virus_2_PRRVS)

AATATGCCAAATAACAACGGCAAGCAGCAGAAGAGAAAGAAGGGGGATGGCCAGCCAGTCAATCAGCTG
TGCCAGATGCTGGGTAAGATCATCGCTCAGCAAAACCAGTCCAGAGGCAAGGGACCGGGAAAGAAAAATA
AGAAGAAAAACCCGGAGAAGCCCCATTTTCCTCTAGCGACTGAAGATGATGTCAGACATCACTTTACCCCTA
GTGAGCGGCAATTGTGTCTGTCGTCAATCCAGACCGCCTTTAATCAAGGCGCTGGGACTTGCACCCTGTCAG
ATTCAGGGAGGATAAGTTACACTGTGGAGTTTAGTTTGCCTACGCATCATACTGTGCGCCTGATCCGCGTCA
CAGCATCACCCTCAGCATGATGGGCTGGCATTCTTGAGGCATCTCAGTGTTTGAATTGGAAGAATGTGTGG
TGAATGGCACTGATTGACATTGTGCCTCTAAGTCACCTATTCAATTAGGGCGACCGTGTGGGGGTGAGATTT
AATTGGCGAGAACCATGCGGCCGAA
